# Delay selection by spike-timing-dependent plasticity shapes efficient networks for signal transmission

**DOI:** 10.1101/2025.10.20.682868

**Authors:** Ali Ghadiri, Hedyeh Rezaei, Ad Aertsen, Arvind Kumar, Alireza Valizadeh

## Abstract

Information processing in the brain relies on efficient communication between different brain regions. Brain oscillations can control signal transmission in brain networks by modulating the timing and excitability of sender and receiver areas. For effective transmission, signals should arrive at target areas when their excitability is maximized. For reciprocally connected neural populations, this mechanism works if the transmission delay matches the period of their evoked oscillation. However, the mechanisms underlying such development of the connections with matched delays remain elusive.

While transmission delays in brain networks change during development, the process by which delays are tuned for efficient transmission is unknown. Here, we demonstrate that the well-known Hebbian learning rule can provide a mechanism for selecting connections with delays that match the period of network oscillations. We consider a reciprocally connected bi-layer network of excitatory and inhibitory neurons that generate network-level oscillations spontaneously or in response to external stimuli. When exposed to spiketiming-dependent plasticity (STDP), the network self-organizes to potentiate connections with delays matching the oscillation period, while depressing those with non-matching delays. Our findings shed light on how transmission delays may evolve during learning and development to optimize the organization of brain networks for efficient signal transmission.

## Introduction

Controllable and flexible communication between different regions along the processing hierarchy and between parallel processing pathways [1–4] is essential for brain function. Collective oscillations of neuronal populations have been proposed as a mechanism for efficient communication between neurons at different spatial scales [3, 5–11]. From a theoretical viewpoint, coordinated neuronal activity through engagement with population oscillations establishes a mechanism for the generation of stronger signals [12, 13], while the change in excitability of the receiver regions over each cycle of the oscillations can determine the effectiveness of the incoming signals [4, 12, 14, 15]. Oscillations can arise as a consequence of the resonance-like response to external stimuli or due to the dynamical state of the brain [16, 17]. When oscillations form a means to facilitate communication between brain networks, it is essential that the oscillations in the sender and receive network are synchronized with a fixed latency. One possible way to synchronize the sender and the receiver network oscillations is to connect the two networks.

However, synaptic connections are always associated with some transmission delays which may depend on the distance between the regions and on the conduction velocity in the neurites that connect the neurons in these regions, both spanning a broad range [18–22].

Key questions are how the brain manages to coordinate spike timing, despite this wide range of delays [20, 23] and whether there exist regulatory processes that may adjust the delays within and between cortical areas [24–29].

Theoretical studies have proposed specific rules for the change in myelination of axons in the central nervous system that might tune the spike arrival time by regulating the transmission delays [30]. However, while there is ample evidence for changes in white matter due to myelination during learning [31], the rules governing this process remain obscure. It is unclear whether myelination is a specific learning process or a homeostatic mechanism for regulating coarse-grained activity, and whether it primarily shapes the structure during development or if it is a lifelong process akin to plasticity [32].

Spike-timing-dependent plasticity (STDP) can also form a mechanism to fine tune delays of the connections by selectively strengthening or weakening connections of specific delays [33, 34]. In this study we propose that the well-known Hebbian STDP rule can shape the distribution of transmission delays for an efficient communication in neuronal networks on its own, without a distinct learning rule acting directly on the transmission delay.

To demonstrate this, we investigated a bi-directional two-layer neuronal network of spiking neurons with inter-layer plastic synapses being updated according to the STDP rule. Our inquiry centered on the behavior of synaptic connections within this framework, particularly the influence of synaptic plasticity on shaping inter-layer connectivity strengths and delays by exploiting the network’s resonance/oscillation property.

Our findings unveil a profound relationship between the network’s oscillation period and synaptic delays of potentiated inter-layer connections when governed by a Hebbian STDP rule. We initiated the model with a wide distribution of delays and found that STDP consistently resulted in the potentiation of synaptic connections whose delays in forward and backward directions added to match the period of oscillations in the two networks. As shown in previous studies, such a match between two seemingly distinct time scales is crucial for efficient transmission of signals along layered neural networks [35, 36]. Thus, we show that through STDP the network self-organizes to a consistent structure where the delays turn out to match the oscillation period, irrespective of whether the oscillations are evoked or spontaneous. This finding suggests a deep interplay between synaptic plasticity and network dynamics, whereby the network fine tunes its temporal properties to optimize signal transmission. In other words, it relates two key properties of the network: resonance/oscillation frequency and the plasticity rule.

## Results

### Baseline activity of the network

Our model consisted of two networks (L1 and L2), each consisting of 200 excitatory and 50 inhibitory neurons. Neurons and synapses parameters in the two networks were identical and therefore the two networks had identical resonance frequencies. Within a network, neurons were recurrently connected (connection probability = 0.2, regardless of the type of neurons). 70 excitatory neurons (projection neurons) from each network projected to the other network forming bi-directional connectivity between the two networks (Fig. 1a).

**Figure 1.**
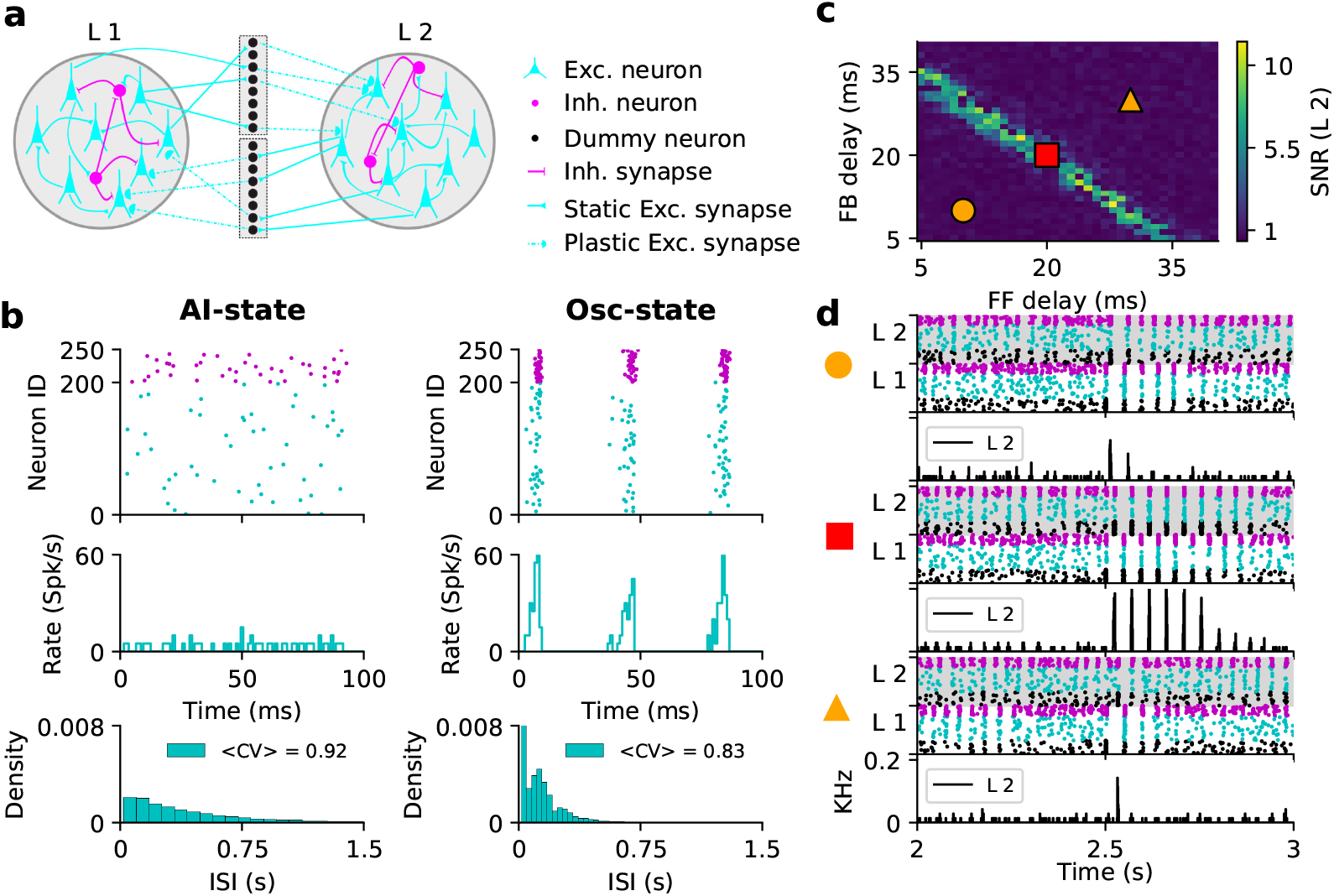
Resonance properties of bidirectionally coupled EI networks. **(a)** Schematic representation of the canonical circuit with a bidirectionally coupled bi-layer network. Each layer contained excitatory (cyan triangles) and inhibitory (magenta circles) neurons sparsely connected by static synapses. Inter-layer connections were excitatory-to-excitatory only, with transmission delays drawn from a normal distribution. FF and FB connections were either static or plastic, depending on the simulation. Axonal delays were modeled using “dummy” neurons (black circles) that transmitted spikes with delay; their number matched the total FF and FB projections. **(b)** Spiking activity of excitatory and inhibitory (E/I) neurons (cyan, magenta; top) and inhibitory synaptic conductance on excitatory neurons (middle) for two input rates producing asynchronous-irregular (AI, left) and oscillatory (right) dynamics. Input rates were 6700 *spikes/s* (left) and 7500 *spikes/s* (right). Bottom: inter-spike interval (ISI) distributions for excitatory neurons. **(c)** FF and FB transmission delays were varied independently to show that matching their total delay to the network’s intrinsic resonance period enables stable pulse packet propagation. **(d)** Spiking activity of two connected layers for three FF-FB delay combinations (Orange circle, red square, orange triangle). Black, cyan, and magenta indicate projection excitatory, excitatory, and inhibitory neurons, respectively.

We tuned the external Poisson input rates to the excitatory and inhibitory neurons to tune the baseline activity of the network in an asynchronous-irregular (AI) state as the baseline state of the network (Fig. 1b, left). Note that the AI state was only approximate, as the spiking activity showed weak signs of regularities (vertical striping in the raster plot (Fig. 1b, left top), which are also reflected in weak oscillations in the averaged inhibitory conductance on the excitatory neurons (Fig. 1b, left middle). The network, however, could also exhibit clear stochastic oscillations. Such a network state could be exposed when the Poisson input rate was increased (input rates provided in the Figure caption) (Fig. 1b, right). For our choice of parameters, the two layer networks had an intrinsic oscillation frequency of ≈25 *Hz*.

Next, we connected the two networks and systematically varied the Feedforward (FF: from L1 to L2) and Feedback (FB: from L2 to L1) delays. For each configuration of FF and FB delays we injected a pulse packet into the sender network (L1) and measured the signal propagation (see Methods). Consistent with our previous work [36], signal propagation (measured by SNR), was maximal when the combined FF and FB delays matched the network’s resonance period (Fig. 1c).

The intrinsic oscillation frequency of our two networks L1 and L2 was ≈25 *Hz*. Hence, signal propagation was most effective when the sum of FF and FB delays approached 40 *ms* (Fig. 1c,d).

### Plastic connections in the bidirectional FF network

To investigate how inter-layer connections evolve under the assigned STDP rule and to clarify the effect of the network’s resonance properties on potentiation and depression of inter-layer connectivity, we connected the two network layers through FF connections. These connections were allowed to change according to the STDP rule. To explore the role of initial delays in determining the polarity (increase or decrease) of synaptic changes, we assumed full connectivity between the excitatory projection neurons in the two populations (*ϵ* = 1). We chose the initial inter-layer delays from a Gaussian distribution (Fig. 2a, blue histograms). We then delivered temporally dispersed pulse packets (number of spikes = 50, sd = 2 *ms*) quasi-periodically (interval = 500 *±* 100 *ms*). The simulation time was 500 *s*, but pulse packets were delivered to the projection neurons in the first layer in times between 1 *s* to 490 *s* (the learning window).

**Figure 2.**
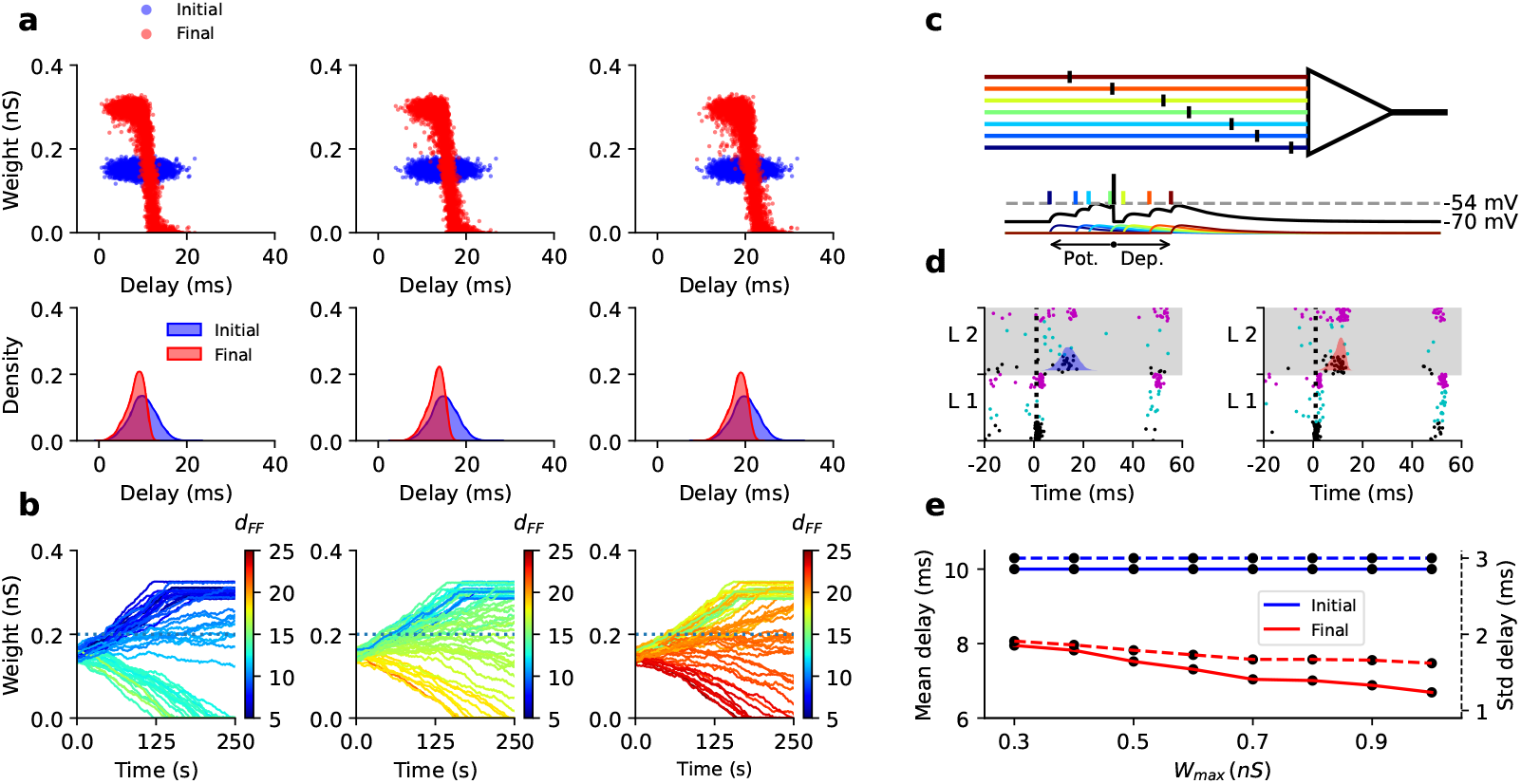
Shorter delays were potentiated in a bidirectional FF network. **(a)** In a bi-layer network with only FF connections, STDP favors connections with shorter delays. Upper panels show scatter plots of synaptic delays and weights at initial (blue) and final (red) states; lower panels show corresponding delay distributions for three initial means (10 *ms*, 15 *ms*, 20 *ms*; *σ* = 3 *ms*). **(b)** Evolution of 50 randomly selected FF connections per condition, color-coded by initial delay, showing potentiation of short-delay and depression of long-delay connections. **(c)** Schematic of how delays influence STDP in a single neuron: short-delay inputs arrive before postsynaptic spikes (pre–post pairing), and hence, give rise to synaptic potentiation, whereas long-delay inputs arrive after (post–pre pairings) resulting in synaptic depression. Colors correspond to delay coding as in **(b). (d)** The bidirectional FF network showed stronger, more temporally focused activation in the second layer and faster spike transmission after learning. Left and right panels show initial and final spiking responses; vertical dashed line marks the pulse packet center of mass. Late spikes vanished as long-delay connections were depressed. With initial FF delays (*µ* = 10 *ms, σ* = 3 *ms*), increasing *W*_*max*_ reduced both the mean (solid) and SD (dashed) of final delays, indicating faster, more reliable transmission. Synaptic weights were drawn from normal distributions: initial *µ* = 0.15 *nS, σ* = 0.01 *nS*; maximum *µ* = 0.3 *nS, σ* = 0.01 *nS*; *W*_*threshold*_ = 0.2 *nS*. Pulse packets were delivered quasi-periodically every 500 *ms* (*±*100 *ms* jitter) over a 500 *s* simulation.

We found that, regardless of the mean of the initial delay distribution, connections with shorter initial delays were potentiated, whereas those with longer initial delays were depotentiated (Fig. 2a). After learning, if the synaptic weight fell below a threshold (horizontal dashed line in Fig. 2b), we considered it as a pruned connection. The distribution of delays of the surviving connections was negatively skewed, with a mean smaller than the mean of the initial distribution (compare red and blue distributions in lower panels of Fig. 2a).

Furthermore, we found that increasing the mean of the initial inter-layer connection delays led to a corresponding increase in the mean of the final delay distribution, though it remained lower than the initial mean in all cases studied (Fig. 2a). That is, connections with shorter delay were more likely to survive and to strengthen (as schematically shown in Fig. 2c). Thus, under the influence of STDP, synaptic weights were not independent of the synaptic delays (Fig. 2a), red dots). Additionally, across all three examples, we observed an increase in the mean of the final synaptic weight distribution, compared to the initial distribution, irrespective of the initial mean delay (Fig. 2a,b).

The reason why connections with shorter delays got potentiation and those with longer delays were depotentiatied is easy to understand: Each neuron in the L2 network received inputs from L1 via connections with distributed delays. If the initial connectivity is strong, inputs with shorter delays may make the postsynaptic neurons spike before the arrival of inputs from the long-delay connections (Fig. 2c,d). Therefore, while the shorter delay connections have pre-post pairing and consequently potentiated, long delay connections effectively end up as post-pre pairing and, hence, get depotentiated.

The potentiation of short-delay projections strengthened the effect of early FF inputs on the second layer, leading to earlier activation of its neurons. As a result, a greater proportion of incoming spikes to the second layer arrived after the post-synaptic spikes, leading to further depression of long-delay connections through the STDP rule. This suggests that increasing the upper limit of synaptic weights causes the depression of more synapses, leading to a narrower final distribution of delays (Fig. 2e).

It should be noted that during the propagation of a pulse packet from L1 to L2, the mechanism underlying the selection of short-delay projections depends on the balance between excitation and inhibition, as well as on the recurrent inhibition in the target layer network (L2). The EI-balance causes the postsynaptic neuron to respond preferentially to the earlier inputs, leading to a potentiation. The synchronous inhibitory activity of L2 rapidly follows the synchronous activity of the excitatory projection neurons of L2 (triggered by pulse packet input from L1) and, thus, significantly reduces the effectiveness of the inputs, leading to a depression. Thus, synapses of postsynaptic neurons that primarily received long-delay projections, arriving after inhibition became dominant, were neither depressed nor potentiated.

### Evolution of FB connections in the bi-layer network

Next, we bidirectionally connected the two layer networks with a connection probability of 0.5. Connection delays were chosen from normal distributions. To characterize the evolution of FB connections, we made the FF connections static, whereas FB connections were allowed to change according to the STDP rule. We then injected pulse packets into the L1 network quasiperiodically (interval = 1000 *±* 200 *ms*) for 1990 seconds. In these simulations, the learning time window was set from 5 *s* to 1995 *s* and simulations were run for 2000 *s*.

First, we chose the FF delays from a Gaussian distribution (mean = 8 *ms* and sd. = 1 *ms*, Fig. 3a, bottom-left, gray distribution). FF weights were drawn from a Gaussian distribution with mean = 0.2 *mS* and sd = 0.01 *ms*. FB connection delays were chosen from a different Gaussian distribution (mean = 25 *ms*, sd = 8 *ms*, Fig. 3a, bottom-left, blue distribution). The initial strength of each FB connection was also drawn from a Gaussian distribution (mean = 0.2 *nS*, sd = 0.01 *ms*, Fig. 3a, top-left, blue distribution). The weights and delays of the FB connections were initially uncorrelated.

**Figure 3.**
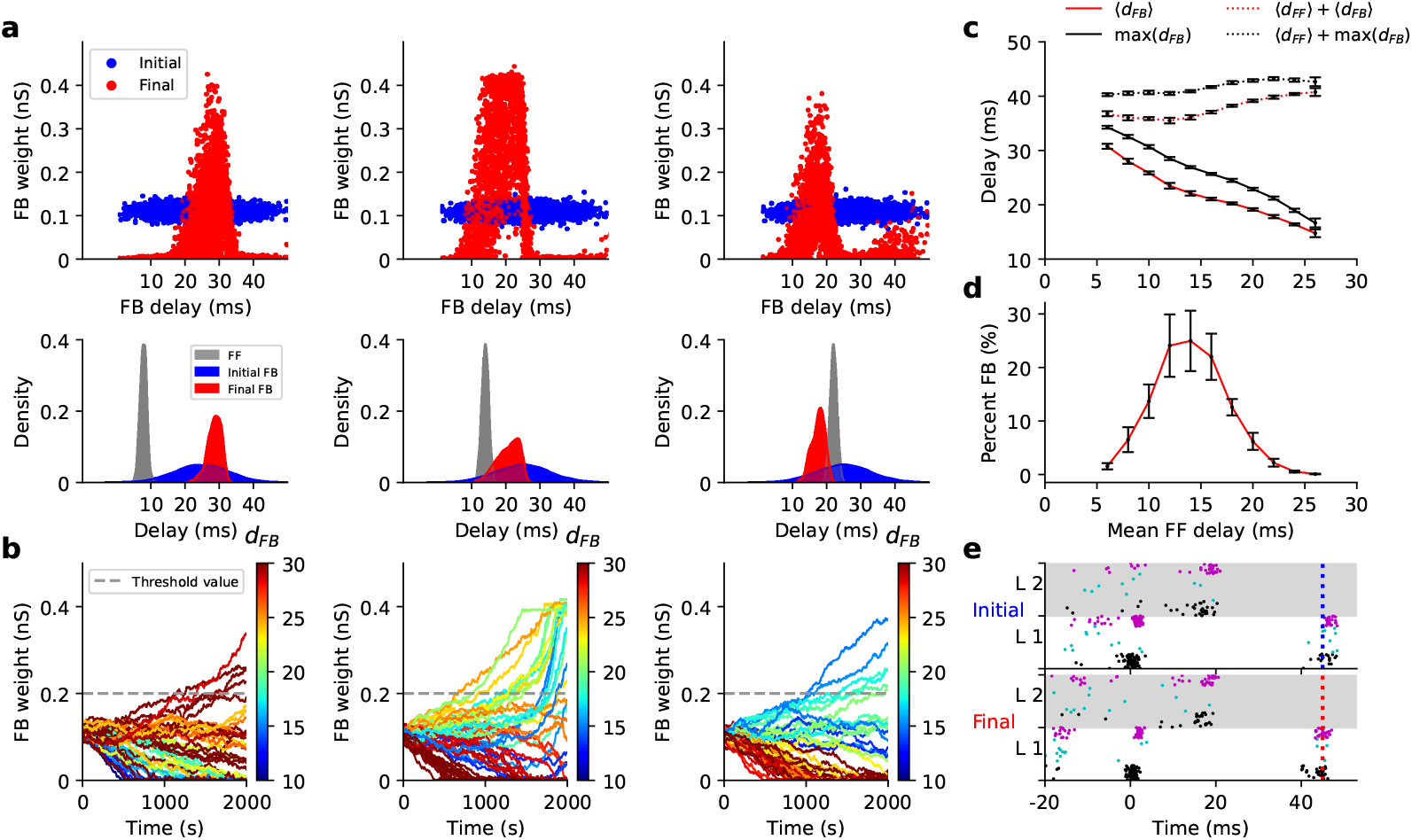
Delay selection of FB connections based on resonance with static FF connections. **(a)** Upper panels show the raster plots of the delays and weights of the FB connections at initial (blue) and final (red) states. Lower panels show the corresponding distribution of the initial delays and final delays of the potentiated FB connections. The FF connections were chosen to be static, with a delay distribution of *σ* = 1 *ms*, and three different mean values (8, 14, and 22 *ms*), as shown in gray in the lower panels. The FB connections evolved through the STDP rule, initialized with a normal delay distribution (*µ* = 25 *ms, σ* = 8 *ms*). As the mean delay of the FF connections increased, the mean of the final distribution of FB delays decreased, resulting in a peak at lower values. **(b)** Evolution of weights for the FB connections with different delays is shown for each set of initial parameters in (a). Legends show the color code used to indicate different delays. **(c)** The mean of the final distribution of FB delays decreased as the mean FF delays systematically increased (red curve). The FB connections evolved such that the final mean delays, summed with the mean FF delays, remained slightly below the both networks’ resonance period. The error bars indicate the standard deviation over the 20 trials. The sum of the maximum values of the final delays in both directions, almost matched the resonance period. **(d)** The number of potentiated FB connections, for different values of the mean delay of FF connections. **(e)** Spiking activity of the two layers at initial and final states. A pulse packet was injected into the first layer network at time 0. FF and FB weights were drawn from normal distributions with parameters (*µ* = 0.2 *nS, σ* = 0.01 *nS*) and (*µ* = 0.11 *nS, σ* = 0.01 *nS*), respectively. The maximum weights allowed also followed a normal distribution (*µ* = 0.4 *nS, σ* = 0.01 *nS*). Additionally, *W*_*threshold*_ was set to 0.2 *nS*. Pulse packets were injected into the L1 network at intervals of 1 *s* with (*±* 200 *ms* jitter) during the period from 5 to 1995 *s*, for a total simulation time of 2000 *s*.

Given repeated exposure to the pulse packet input, FB connections changed over time (Fig. 3b left). As expected, synaptic changes depended on their respective connections delays. In this example, most potentiated connections lay in a small range of delays (25 − 35 *ms*). Outside this range, all synapses were depotentiated (Fig. 3a, top left, red histogram). Clearly, STDP created a correlation between synaptic weights and delays. Similar to the previous Section, we disregarded all connections with weight smaller than a threshold (0.2, Fig. 3b), and observed that the resulting delay distribution had a mean delay of ≈30 *ms*.

Similar results were obtained when we started the simulations with larger mean delays in the FF connections (Fig. 3a, bottom, middle and right panels). However, as we increased the mean delay of the FF connections, the mean delay of the surviving FB connections decreased. In all cases, however, the sum of the maximum delay of the potentiated synapses in the FB direction with the maximum forward delay remained close to the resonance period (Fig. 3b,c). In Fig. 3c, we demonstrate that not only the sum of the maximums of FF and survived FB delays aligns with the network’s resonance period (black dotted line), but also that the sum of their mean values shows a close correspondence (red dotted line). Nevertheless, the match is even more precise in the former case.

As expected, the number of surviving (potentiated) FB connections depended on the presence of FF connections with matching delays in the initial distribution. Fig. 3d shows the number of potentiated FB connections as the mean of the initial FF delay distribution was systematically varied. This result indicates that the number of potentiated FB connections peaked when the sum of the mean FF delay and the mean initial FB delay matched the resonance period of the two layer networks.

Applying the STDP rule to FB connections enabled the bi-layer network to approach resonance. Initially, the broad range of delays in the feedback pathway caused backward-propagating spikes to be temporally dispersed. However, after STDP induced a selection of suitable FB connection delays, spikes from network L2 arrived in network L1 much more synchronously (Fig. 3e).

### Self-organization of the delays when both FF and FB connections were plastic

We next made both FF and FB connections plastic. In the initial state, both FF and FB connection weights and delays were drawn from a Gaussian distribution, with the initial FF delay distribution being narrower than the initial FB delay distribution (Fig. 4a, left). Note that as in the previous simulations, initial synapse weights and delays were not correlated. To train the FF and FB connections, we injected pulse packets in a quasi-periodic manner (interval = 1000 *±* 200 *ms*). In these simulations, pulse packets were injected into the first layer network only between 5 *s* to 2495 *s* (learning time window) and the networks were simulated for a total of 2500 *s*.

**Figure 4.**
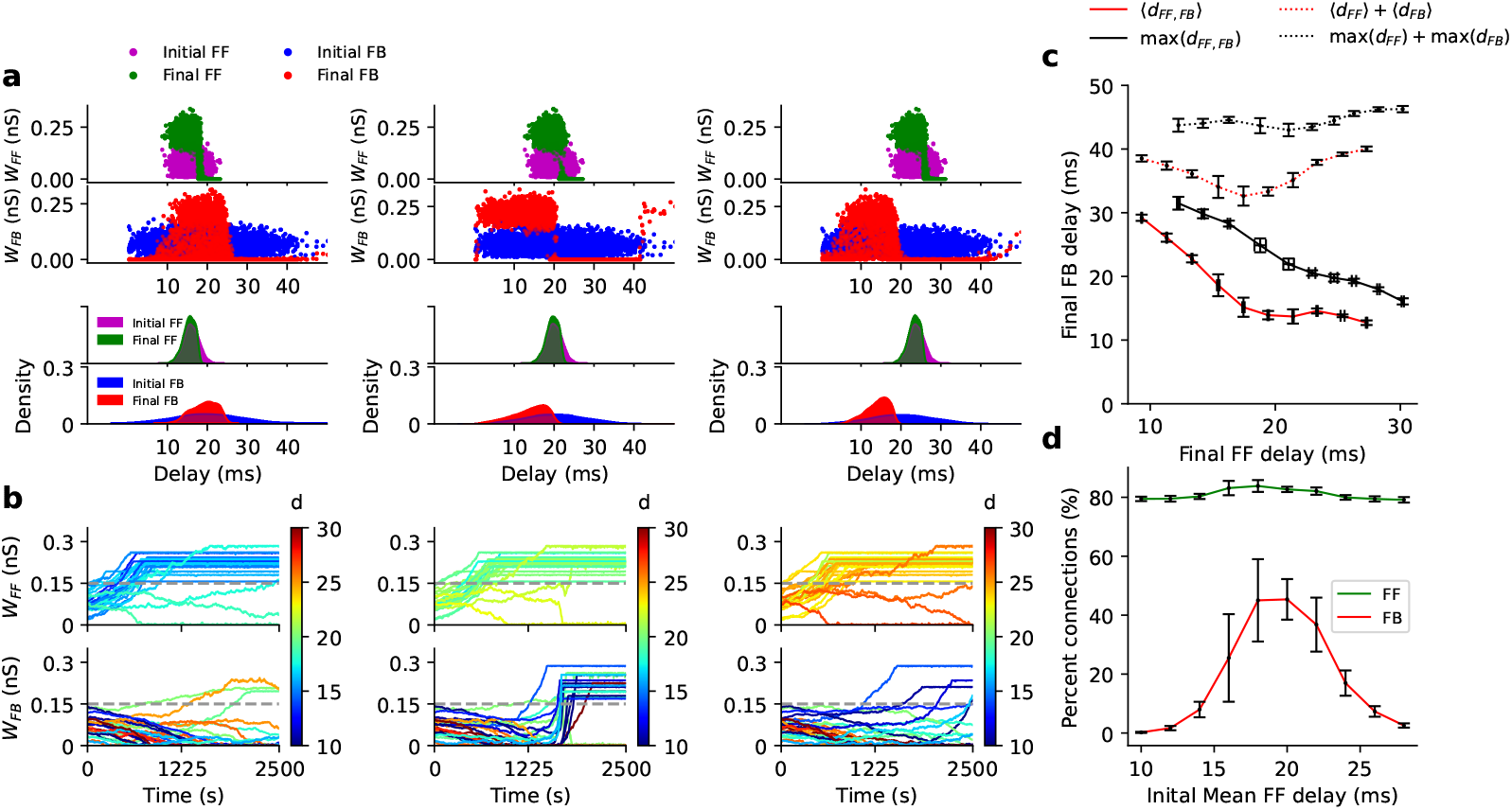
Delay selection in both directions based on resonance. **(a)** Raster plots of the initial and final values of delays and weights of FF and FB connections (upper panels) and the initial and final distributions of FF and FB delays are shown for three different initial mean FF delays (16 *ms*, 20 *ms*, and 24 *ms*) with *σ* = 2 *ms* (top panels; scatter plots). The initial FB delays followed a normal distribution (*µ* = 20 *ms, σ* = 8 *ms*). **(b)** The evolution of 20 randomly selected FF and FB connections for mean FF delay of 15 *ms*, are shown, revealing faster modification of FF connections, compared to FB connections. **(c)** As the mean of the FF delays increased, the mean and maximum value of the final FB delays (red and black solid curves) decreased, so that their sum (dotted lines) remained approximately constant and close to 40 *ms*, the resonance period of each layer network. **(d)** Number of the FF and FB connection in the final state, for different values of mean initial FF delay. The initial FF and FB weights were drawn from a normal distributions (*µ* = 0.07 *nS, σ* = 0.03 *nS*). The maximum weight allowed through the STDP rule, followed a normal distribution (*µ* = 0.22 *nS, σ* = 0.03 *nS*) with a threshold set to *W*_*threshold*_ = 0.15 *nS*. Pulse packets arrived at intervals of 1 *s* (with *±* 200 *ms* jitter) within the time window of 5 to 2495 *s*, for a total simulation duration of 2500 *s*. All panels show results averaged across 20 trials.

As in the previous simulations, STDP potentiated synapses with specific delays and, thereby, created a correlation between synaptic weights and delays for both the FF and FB connections (Fig. 4b, left). We found that FF connections consistently maintained shorter delays in the final distribution. This indicates that, regardless of the networks’ resonance properties, FF connections with longer delays were pruned during the transmission of pulse packets in the forward direction (Fig. 4a).

By contrast, FB connections were tuned according to the resonance properties of the individual networks. A change in the mean of the FF delay distribution also resulted in a corresponding change in the FB delay distribution: increasing the mean FF delay resulted in a decrease in the mean final FB delay (Fig. 4a,c). Across all conditions, the sum of the mean (as well as the sum of the maximums of) final delays for FF and FB connections remained relatively close to the intrinsic oscillation period (respectively, red and black dotted curves in Fig. 4c).

Importantly, the emergence of FB connections with delays that matched the networks’ resonance properties depended on the presence of a sufficient number of connections with appropriate delays in the initial FB delay distribution. The mean of the FF delays determined the range of delays that were effectively potentiated through plasticity. Therefore, the number of FB connections that were potentiated depended on the mean FF initial delay (Fig. 4d).

Note that compared to the case of static FF synapses (Fig. 3d), the peak was moved to even shorter FF delays, since here the mean final FF delay itself was shorter than the initial one.

Because FB connection weights and delays depend on the FF delay distribution to favor appropriate delays, FB connections reached their steady state later than the FF connections (Fig. 4b).

Tuning of FF and FB synapses using STDP led to an improved propagation of signals from L1 to L2. Fig. 5 shows that the pulse packets initially failed to reverberate in the network, whereas after synaptic modification via STDP, successful reverberation between the layer networks was established. It is worth noting that the facilitation of signal propagation, quantified by an increase in the signal-to-noise ratio, SNR, did not trivially come with an increase in synaptic cost, defined as the sum of all synaptic weights of FF and FB connections (Fig. 5). While the final synaptic cost depended on the initial distribution of delays and weights of the inter-layer connections, our results suggest that improved signal transmission is not merely a trivial outcome of increased mean connectivity weight between the two layers.

**Figure 5.**
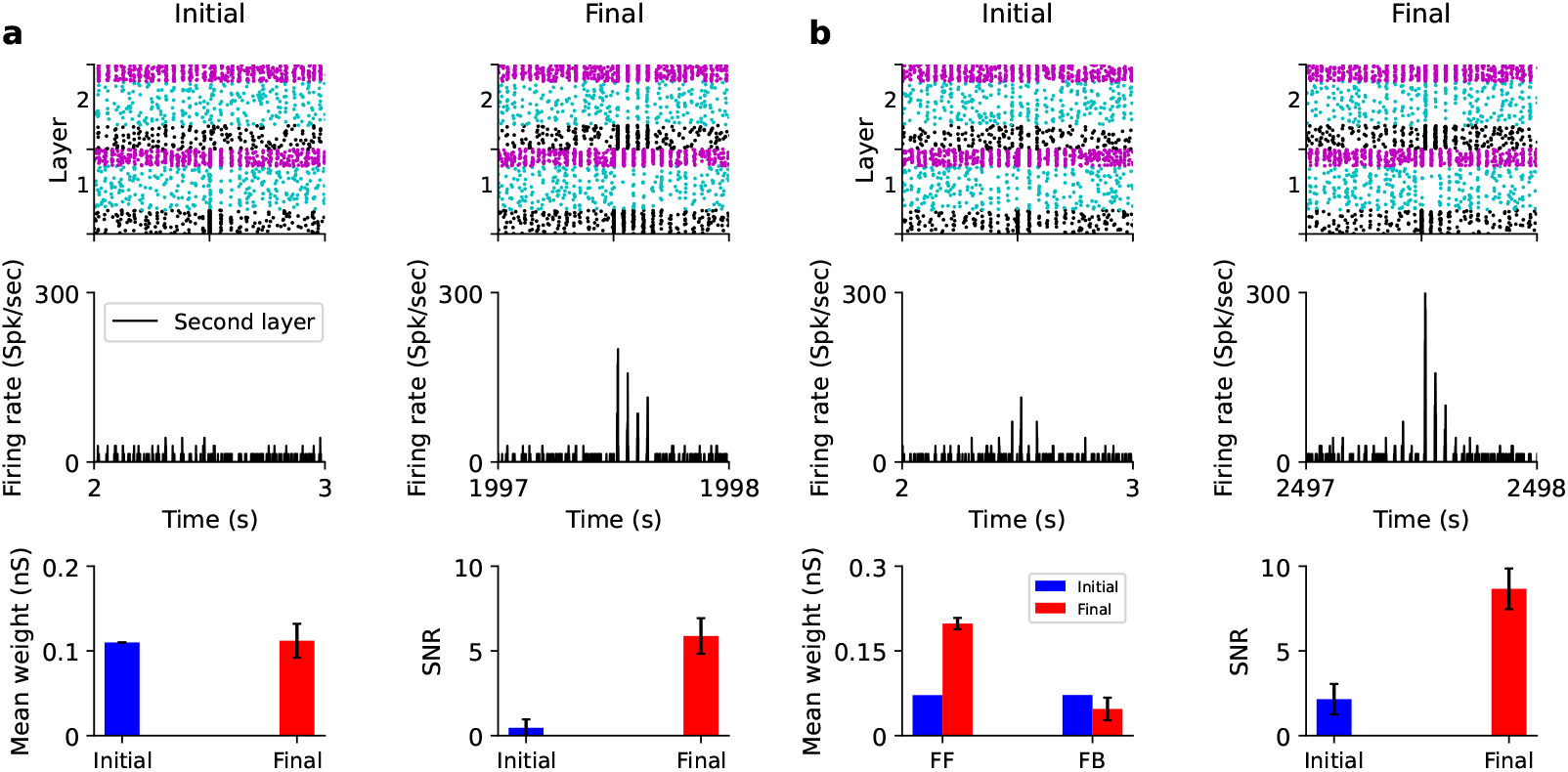
STDP facilitates pulse packet propagation through delay selection. A bi-layer network successfully transmitted a single pulse packet (*α* = 40, *σ* = 2 *ms*) after modification of connectivity under two distinct configurations: static FF and plastic FB connections (a), and both plastic FF and FB connections (b). In both configurations, the second layer exhibited a high SNR due to the alignment of inter-layer delays with the resonance periods of the individual layer networks. (**a, b**) By stimulating the first layer with a single pulse packet, we recorded the spiking response in the second layer. After synaptic modification through STDP, the pulse packet successfully reverberated between the two layer networks. Also, the higher value of SNR did not rely on excessive values of synaptic strengths, as the mean weight of the resonant connections might keep their mean final weights around or even below the initial ones (bottom panels). The parameters used in (a) and (b) were the same as those in Figs. 3 and 4, respectively. The error bars indicate the standard deviation over 20 trials.

### Delay selection in the oscillatory regime

Thus far, we have presented pulse packets quasi-periodically to train the synapses. Such a training could be considered non-biological. To alleviate this problem, we hypothesized that spontaneous oscillations in the dynamics of neuronal networks can facilitate the formation of the connections in a specific range of delays. That is, oscillatory activity can provide pulse-packet-like input to the network and train the synapses. To test this hypothesis, we set the parameters of the two networks such that they operated in an oscillatory regime (Osc state; Fig. 1b, right column) and the population activity showed oscillations at ≈25 *Hz*.

Initially, we considered unidirectional FF connectivity between two identical layer networks, characterized by a delay distribution with an initial standard deviation of 3 *ms*, and let them evolve through the assigned STDP rule. We found that, regardless of the initial mean delay, connections with shorter delays were preferentially potentiated (Fig. 6a, left panel). As a result, both the mean and the standard deviation of the delay distribution decreased in the final state. Furthermore, increasing *W*_*max*_ led to a progressive reduction in both the mean and the width of the final delay distribution (Fig. 6a, right panel), resembling the behavior observed in the case of evoked activity discussed earlier (Fig. 2d). Notably, these simulations predict that, in spontaneously oscillatory networks, the oscillatory activity of the L2 population would become phase-locked to that of the L1 population, with a time lag approximately equal to the mean delay of the inter-layer connections. Consistent with this hypothesis, the time lag between L1 and L2 population activity scaled with the mean of the initial delay distribution (Fig. 6b, right panel). Three examples of phase-locked spiking activity of the two layer networks are represented for the three different values of FF delays (Fig. 6b, left panel).

**Figure 6.**
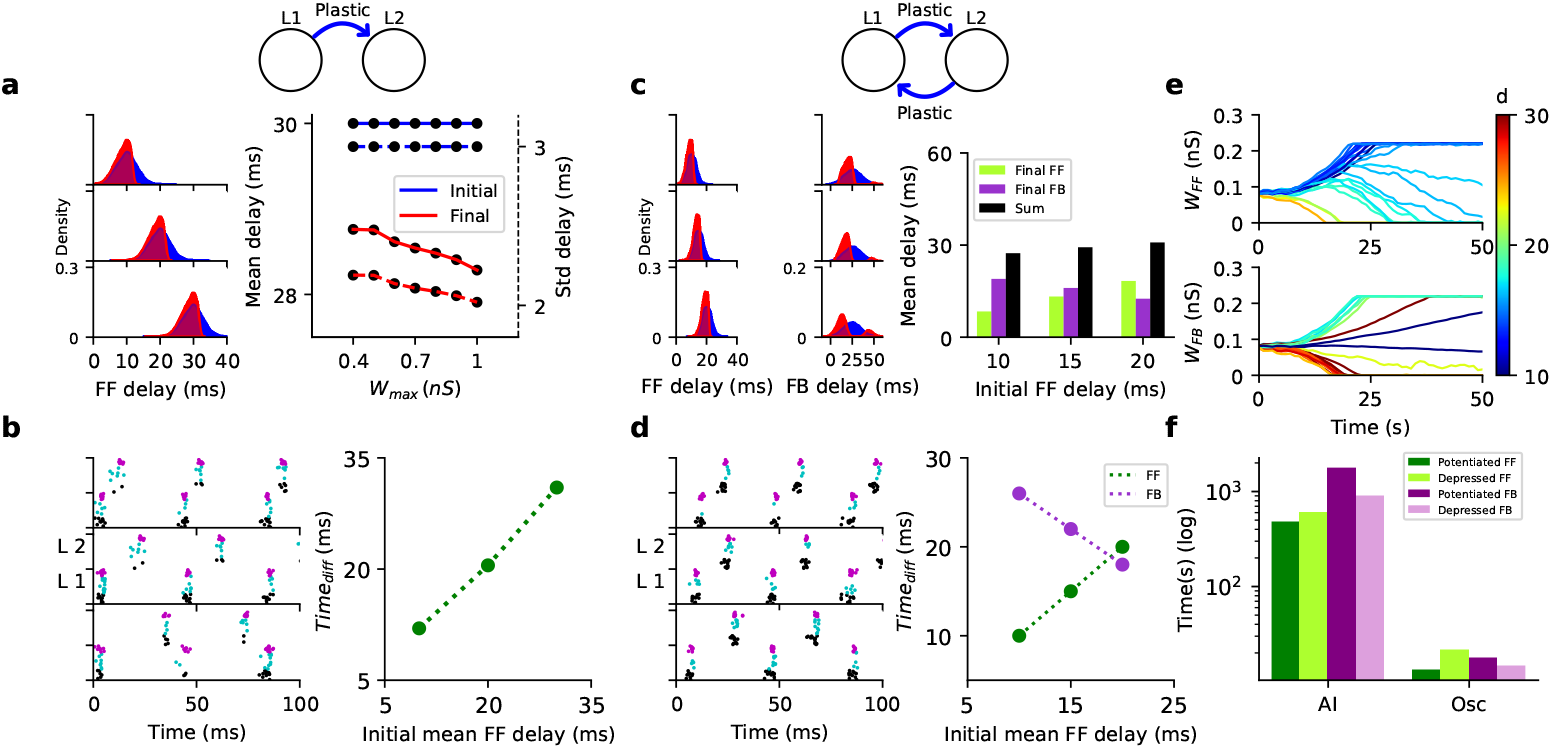
Delay selection in unidirectional and bidirectional oscillatory layer networks. **(a)** In a bi-layer oscillatory network with unidirectional projections, STDP favored short-delay connections for all initial mean delays. Left: initial (blue) and final (red) interlayer delay distributions after 30 *s* of learning. Right: increasing *W*_*max*_ reduced the mean and SD of final delays, indicating more depressed long-delay connections. Initial mean FF weight was 0.2 *nS*; *W*_*max*_ = 0.4 *nS, W*_*threshold*_ = 0.2 *nS*. **(b)** Spiking activity for each initial condition during 100 *ms* after 26 *s* of learning. Right: relation between inter-layer spike timing difference and initial mean FF delay. Spike times were computed as the center of mass of synchronized volleys, averaged across phase-locked events within 1 *s* after 26 *s* of learning. **(c)** In bidirectionally connected oscillatory networks, both FF and FB connections evolved via STDP. For three initial FF delay means (*µ* = 10, 15, 20 *ms, σ* = 3 *ms*), final delay distributions after 50 *s* (left; blue: initial, red: final) showed that total loop delays (black bars) remained constant (right). FB delays had *µ* = 25 *ms, σ* = 10 *ms*. Initial inter-layer weights: 0.08 *nS*; *W*_*max*_ = 0.22 *nS*; *W*_*threshold*_ = 0.15 *nS*. **(d)** Spiking activity during 100 *ms* after 46 *s* of learning (left) and corresponding inter-layer timing differences (right), as in (b). **(e)** Evolution of 40 randomly selected FF (top) and FB (bottom) connections for initial mean FF delay of 15 *ms*. **(f)** Connectivity over-time evolution—both potentiation and depression—occurred faster in the oscillatory state (panel **(e)**) than in the stimulus-evoked AI state (Fig. 4b). Bars show the time for each connection type to cross the weight threshold after potentiation (0.15 *nS*) or depression (0.03 *nS*).

We next considered the reciprocally connected network. To break the network’s symmetry and identify up- and down-stream layers, we initialized the FF connection delays using a relatively narrow distribution. By contrast, FB connections were initialized with a wider delay distribution, consistent with the case of evoked activity discussed earlier (Fig. 4).

We observed that the evolution of the connections ensured that the sum of the final mean delays in both directions remained approximately constant, aligning with the period of the oscillatory activity (Fig. 6c). Furthermore, the means of the final delay distributions determined the phase relationship between the oscillations of the bi-layer networks. Consistent with the emergent delay distributions, the phase difference between the oscillations of the two layer networks corresponded to the delays in both directions. As shown in Fig. 6d (left panel), the time difference in spiking activity between the first and second layer networks aligned closely with the mean of the delay distribution in each configuration.

A key observation regarding the evolution of inter-layer connectivity in the spontaneous oscillatory state is that the formation and stabilization of consistent delays occurred significantly faster than in the stimulus-evoked activity in the AI state studied earlier (Fig. 6f, compare to Fig. 4). This difference is reasonable since, in the stimulus-evoked activity in the AI state, effective weight modification happened only after the impact of each input pulse packet. By contrast, in the oscillatory regime, weights were updated at each oscillation cycle, which is much shorter than the intervals between stimulus consecutive pulse packets in the stimulus-evoked AI state, which was set to be around 1 *s*.

## Discussion

The adaptability of signal transmission speed and synaptic delays in the brain has been observed in experimental studies, suggesting that delays are not static, but can change dynamically in response to neuronal activity [37–42]. However, the specific rules governing this delay plasticity remain largely unknown. Computational studies have proposed mechanisms in which delay plasticity operates in parallel with synaptic weight plasticity, demonstrating that such dual plasticity can enhance the computational capabilities of neuronal networks [43, 44]. In our study, we show that synaptic delays can be tuned without requiring a specific mechanism for delay plasticity. Instead, we demonstrated that the STDP rule, traditionally associated with synaptic weight changes, is sufficient to modify delays implicitly and effectively.

By allowing a STDP rule to shape synaptic connections based on precise spike timing, delays naturally evolve to align with the networks’ resonance properties, optimizing communication efficiency. This approach builds on earlier work [33, 45], which showed how STDP could refine temporal relationships between neurons’ spiking activities. Our findings highlight an additional dimension, though: the ability of STDP to exploit network oscillatory dynamics to fine tune delays in reciprocally interconnected neuronal networks. This integration of resonance properties into the delay selection process adds another aspect of functional significance, demonstrating how a single plasticity mechanism can simultaneously coordinate synaptic weights and delays to achieve optimal signal propagation and network synchronization.

We speculated that these findings could be of importance for early developmental processes, during which neural circuits undergo extensive refinement driven by spontaneous and evoked activity [46]. Previous research suggested that early endogenously generated patterns of activity play a crucial role in guiding the formation of neural connections in sensory and motor areas [47–50]. Our results support this hypothesis by showing that synaptic reorganization induced by spontaneous oscillatory activity was compatible with that of stimulus-evoked activity, whereas the modification through the endogenous rhythmic activity was considerably faster than that induced by the evoked activity. The continuous nature of spontaneous oscillations enabled rapid weight updates within each oscillation cycle, aligning delays more rapidly and efficiently. This finding underscored the developmental importance of oscillatory rhythms in shaping the temporal structure of neuronal networks along with the sensory-driven activity.

Moreover, our results suggest that oscillations can stabilize the tuning of delays beyond the brain development period. Disruptions in oscillatory dynamics and timing mechanisms have been implicated in several neurodevelopmental disorders, including autism, schizophrenia, and epilepsy [51]. Our findings suggest that aberrant oscillatory activity during development and beyond could impede the proper tuning of inter-network delays, leading to inefficient communication and cognitive deficits. Investigating how oscillatory disruptions affect delay tuning in pathological states could provide new insights into the etiology of these conditions and suggest novel therapeutic strategies.

How fast the networks will develop their inter-connectivity well enough to facilitate communication in the resulting circuitry depends on the shape of the STDP kernel and the oscillation frequency of the networks involved. STDP seemed to be most effective in changing synapses in the *θ* band (6 − 10 *Hz*) as in that range consecutive spikes are far apart to exploit pre-post pairing, while minimizing the post-pre interactions [52]. However, resonance in the *θ* band will considerably slow down the transmission rate (which depends on the resonance frequency) [53]. One possibility might be to train the FF and FB connections by external oscillatory inputs.

Connections between layer networks should be flexible enough, so that they can be modified in a context/state dependent manner. Here we have described a strategy by which connections with certain delays were selected to facilitate communication between two layer networks. How could we also unlearn these connections? The simplest way to weaken the connections in the model would be to alter the resonance properties of the receiver network, which would entail a change in the excitation-inhibition balance and the gain of individual neurons [53]. In addition, a change in the spiking mode of neurons, for example a change from single spike emission to bursting behavior, could also alter the oscillatory dynamics and, thereby, weaken the synapses [54].

In general, connectivity between two network operating in oscillatory regime depends on both frequency and phase of oscillations [55]. In fact, interaction between STDP and oscillations can make the network structure bistable and depending on the initial conditions, there can be no connectivity, unidirectional connectivity or bidirectional connectivity. In our model, delays introduced phase differences between the two network oscillations and therefore connections corresponding to specific delays survived. However, if some process were to change the phase of oscillations (e.g. inputs from external sources) the network connectivity could be completely reorganized [55].

### Model Limitations and Future Directions

While our study provided valuable new insights, it simplified certain aspects of cortical circuits. We used a conventional linear profile of STDP [56–58], to highlight its role in the selection of proper connection delays and synaptic weights. Several theoretical and experimental studies have shown the shortcomings of this type of STDP rule, and have suggested variants and complementary mechanisms [43, 59–65]. Moreover, we assumed homogeneity in STDP rules, whereas biological networks exhibit diverse forms of plasticity that may influence delay tuning [66]. Incorporating heterogeneous plasticity rules could refine our understanding of how delays are tuned across different brain areas and oscillatory regimes.

The model networks we studied consisted of several hundreds of neurons. The question arises, whether the phenomena of delay tuning via STDP and its interactions with network dynamics also hold for much larger networks as we have them in real brains. As we have shown, the key observation of delay tuning is a property of network oscillations. Therefore, as long as a network is capable of network oscillation and resonance properties, we should observe this kind of delay tuning, independent of the network size, provided the scaling of the various parameters is suitably taken into account. Moreover, both small and larger networks can show oscillations, as we know from many studies in the experimental and theoretical literature.

Another limitation is the assumption of static resonance properties within the coupled networks. In biological systems, resonance frequencies are dynamic and can be modulated by neuromodulators, learning, or disease states [67]. Exploring how such modulation influences delay tuning would deepen our understanding of the adaptability of neural circuits. Finally, experimental validation of our findings remains a challenge. Advances in high-resolution imaging and optogenetics are essential for directly measuring the evolution of synaptic delays and resonance properties *in vivo*.

Despite its simplifications, the model provides a robust framework for understanding the interplay between synaptic plasticity, connection delays, and oscillatory network dynamics.

Insights from this work open new avenues for investigating how delay tuning mechanisms may contribute to cognitive function and dysfunction.

## Data Availability

The corresponding codes will be deposited in a Github repository and made available upon publication.

## Methods

### Neuron and synapse model

The neuron model simulates a spiking neuron by employing integrate-and-fire dynamics (Equation 1) using conductance-based synapses:

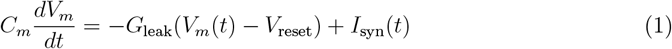

where *V*_*m*_ denotes the membrane potential, *C*_*m*_ the membrane capacitance, *G*_*leak*_ the membrane leak conductance, *V*_*reset*_ the reset potential after initiating a spike, and *I*_*syn*_ the total synaptic input current.

When the membrane voltage reached the threshold for spiking (*V*_*th*_ = − 54 *mV*), a spike was emitted, and the membrane potential was reset and clamped to the resting membrane potential (*V*_*reset*_ = − 70 *mV*) for a refractory period of *τ*_*ref*_ = 2 *ms*.

To avoid a transient network synchrony at the beginning of the simulation, initial membrane potentials of neurons were drawn from a uniform distribution (low: − 70 *mV*; high: − 60 *mV*). Neuron model parameters are listed in Table 1.

**Table 1:** Summary of Model Parameters.

| Name | Value | Description | Name | Value | Description |
| --- | --- | --- | --- | --- | --- |
| **Neuron model parameters** |  |  |  |  |  |
| $C_m$ | 250 pF | Membrane capacitance | $G_{\text{leak}}$ | 16.67 nS | Membrane leak |
| $V_{\text{th}}$ | -54 mV | Firing threshold | $V_{\text{reset}}$ | -70 mV | Reset potential |
| $\tau_{\text{ref}}$ | 2 ms | Refractory period | | | |
| **Synapse model parameters** |  |  |  |  |  |
| $\tau_{\text{exc}}$ | 3 ms | Excitatory decay | $\tau_{\text{inh}}$ | 8 ms | Inhibitory decay |
| $E_{\text{syn}}^{\text{exc}}$ | 0 mV | Excitatory reversal | $E_{\text{syn}}^{\text{inh}}$ | -80 mV | Inhibitory reversal |
| $J_{ee}$ | 0.33 nS | E→E | $J_{ei}$ | 5.5 nS | E→I |
| $J_{ie}$ | -6.2 nS | I→E | $J_{ii}$ | -15 nS | I→I |
| $J_{pe}$ | 0.25 nS | Poisson→E | $J_{pi}$ | 0.4 nS | Poisson→I |
| $J_{pp}$ | 0.6 nS | Pulse Packet→P | $d_E$ | 1 ms | Excitatory delay |
| $d_I$ | 2 ms | Inhibitory delay | | | |
| **STDP parameters** |  |  |  |  |  |
| $\lambda$ | 0.005 | Learning rate | $\alpha$ | 0.9 | Depress./potent. scaling |
| $\tau_+$ | 15.9 ms | Potentiation | $\tau_-$ | 19.3 ms | Depression |
| **Network model parameters** |  |  |  |  |  |
| $N_{\text{exc}}$ | 200 | Excitatory neurons | $N_{\text{inh}}$ | 50 | Inhibitory neurons |
| $N_{\text{proj}}$ | 70 | Projecting neurons | $\epsilon$ | 0.2 | Within-layer prob. |
| $\epsilon_{pp}$ | 1 | Inter-layer prob. | | | |

Synaptic inputs from within-layer connections were introduced through a transient modification of the synaptic conductance *G*_*syn*_:

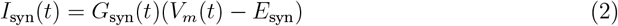

where *E*_*syn*_ is the synaptic reversal potential. Conductance changes were modeled as single exponential functions:

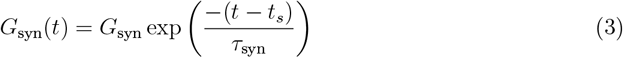

where *G*_*syn*_ denotes the connection strength, and *τ*_*syn*_ is the synaptic time constant. This function is normalized to ensure that a spike event with a weight of 1.0 corresponds to a peak conductance of 1 *nS*. Synapse model parameters are given in Table 1. Synaptic transmission delays were set to 1 *ms* (excitation) and 2 *ms* (inhibition) for within-layer connections, whereas inter-layer feedforward (FF) and feedback (FB) transmission delays were drawn from normal distributions, the parameters of which (mean and standard deviation) were varied as mentioned in each Figure caption.

Inter-layer connections were among excitatory neurons in the two successive layer networks. In the case of static connections, inter-layer connections (for FF connections only), the synapses were as strong and as probable as within-layer excitatory to excitatory connections. For plastic inter-layer connections (involving FF or FB connections as mentioned in each Figure caption), though, synapse weights were subjected to a plasticity rule.

Synaptic weights of plastic inter-layer connections varied according to a temporally asymmetric Hebbian STDP rule as follows:

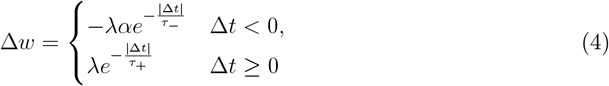

The STDP parameters are given in Table 1. Here, we treated the case where all spike pairs contributed to weight changes. Depending on the relative timing of the spike pairs, the plasticity rule led to potentiation or depression, respectively. The STDP rule applied in this manuscript was weight-independent, by which a bimodal distribution for final weights was achieved. This, in fact, was a positive mark, because two classes of synapses were distinguishable: very weak and very strong connections. Therefore, synapses that were stronger than a well-defined strength threshold (*W*_*threshold*_) were considered potentiated and contributed to the final distribution of delays. Otherwise, they were considered depressed and played a less or no important role in the final distribution and, hence,in signal propagation. However, as described, a hard upper boundary was required to avoid instabilities in the network dynamics due to over-potentiation of inter-layer connections.

Instead, a weight-dependent STDP rule, in general, takes care of this kind of instability by incorporating the current weights in weight updates. However, it hardly takes into account clearly differentiated depressed and potentiated synapses. Hence, we applied a weight-independent STDP rule with a hard upper boundary (*W*_*max*_) in our layered network to study the final distribution of potentiated inter-layer delays and investigate their role in signal propagation across the network.

## Network connectivity

The network consisted of two layers, each comprising 200 excitatory and 50 inhibitory neurons (Fig. 1a). Within a later 70 neurons projected to the other layer. Random connectivity within the layers was established with a fixed connection probability of 0.2 for all types of connections (EE, EI, IE, and II). The intra-layer synaptic weights were kept constant at the values specified in Table 1.

Initially, the study focused on the Asynchronous-Irregular (AI, [68]) regime, characterized by network-level resonance properties, i.e., the appearance of damped oscillations with a certain resonance frequency in response to external synchronous spike volley input (so-called **pulse packets** [69, 70]). Subsequently, the investigation shifted to the Oscillatory (Osc) regime, characterized by the emergence of spontaneous oscillations in the network activity.

Within each regime, both static and plastic inter-layer connections were considered. We examined three scenarios for the modification of inter-layer connections: (i) FF plastic connections in a FF bi-layer network, (ii) static FF and plastic FB connections in a reciprocally connected network, and (iii) plastic FF and FB connections in a reciprocally connected network. In each scenario, the initial strengths and delays of FF and FB connections were drawn independently from normal distributions. For plastic connections, the initial weights and delays were determined, based on assigned distributions, with the potential to change via the STDP rule. By contrast, static inter-layer connections retained their weights and delays as per the assigned distributions.

Inter-layer connections were exclusively excitatory. In experiments with static inter-layer connectivity, the connection probability was set to 0.5, involving 70 projection neurons per layer. As a result, each projection neuron in the second layer received 40 excitatory inputs from within its own layer and 35 excitatory inputs from the other layer, on average. For all experiments involving plastic inter-layer connections, the initial connectivity between the two sub-populations of projection neurons was set all-to-all.

Self-synapses and multiple synapses were not included in the network configuration. Additional details on the network parameters are given in Table 1.

### External input

To achieve the asynchronous irregular network state indicated in Fig. 1b (left column), each excitatory neuron in each layer was driven by 6, 700 independent Poisson excitatory spike trains, each with a mean rate of 1 *spike/sec*. Also, each inhibitory neuron in each layer was driven by 4, 000 independent Poisson excitatory spike trains, each with the same mean rate. However, for the oscillatory state represented in Fig. 1b (right column), we changed the input to the excitatory nwurons in each layer to 7, 500 independent Poisson excitatory spike trains, again each with a mean rate of 1 *spike/sec*. The input to the inhibitory neurons in each layer remained the same. These input parameters, though, were systematically varied in Fig. 2a to investigate the state space of possible network activity regimes.

The external pulse packet stimulus was injected to projecting neurons in the first layer. Each pulse packet consisted of a fixed number of spikes (*a* = 50), randomly distributed around the arrival time (*t*_*n*_). The time of each individual spike was drawn independently from a Gaussian distribution centered around *t*_*n*_, with a standard deviation of *σ* = 2 *ms*.

## Data analysis

### Population Fano Factor

To differentiate the dynamical states of the network, we used the population Fano factor (pFf). Higher values of pFf indicate more synchronized neuronal activity, whereas lower values denote a more irregular activity regime. To estimate the population Fano factor, we divided the time interval into bins of size Δ*t* = 5 *ms* and transformed the population spike trains into spike count vectors *y*_*i*_(*t*) using a rectangular kernel. The population Fano factor was then calculated as:

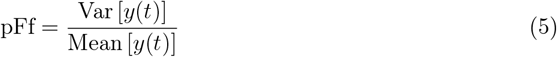

### Signal to Noise Ratio

To distinguish successful propagation of single input pulse packets from failed propagation, we estimated the Signal-to-Noise Ratio (SNR). This measures the ratio of the variance of the averaged membrane potential of neurons in the second layer network (the

receiver network) upon pulse packet injection into the first layer network, normalized by its variance during ongoing network activity:

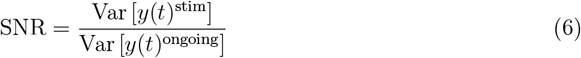

### Power Spectral Density Analysis

The spiking times for each layer were converted into a spike count vector with a bin size of 1 *ms*. To analyze the frequency content of spiking activity over time in the oscillatory network state, we employed Welch’s method, a widely used technique for estimating the Power Spectral Density (PSD). This method partitions the spike count data into overlapping windows, each with a 50% overlap between consecutive segments ([0 *s*, 10 *s*], [5 *s*, 15 *s*], …). The PSD was then computed for each window segment.

## Simulation tools

Network simulations were performed using the simulation tool NEST (www.nest-initiative.org) [71–73], interfaced with PyNest. The differential equations were integrated using fourth order Runga-Kutta with a time step of 0.1 *ms*.

In NEST, the delay implemented in connection models corresponds to dendritic delay. To introduce an additional axonal delay, which was fundamental to our study, two intermediate so-called “dummy neuron” populations were included for both forward and backward directions (Fig. 1a). These dummy-neuron populations were configured to introduce the desired axonal delay required for the specific simulation. For each inter-layer connection, one dummy neuron was considered, such that any dummy neuron had only one input and one output connection. Connections from the presynaptic neuron to the dummy neuron were strong, so that each dummy neuron was activated following each spike from the presynaptic neuron connected to it, after a time delay that was the desired axonal delay in the experiment. The connection strength from dummy neurons to postsynaptic neurons was equal to the corresponding biological synapse. Connections from dummy neurons to target neurons were subjected to the plasticity rule. Additionally, dendritic delays were implemented in these connections. In other words, the output connection of each dummy neuron targeted a postsynaptic neuron, included a dendritic delay, and its synaptic strength was equal to the strength of the actual connection between the pre- and postsynaptic neurons. Then, this synapse could be modified through the STDP rule (Fig. 1a).

It is worth noting that delays could be honored on both incoming and outgoing connections from and to dummy neurons. We assigned the fixed value of 0.1 *ms* to the dendritic delays, whereas the axonal delays (those of connections between presynaptic neurons and dummy neurons) were drawn from a normal distribution with mean and standard deviations indicated in each Figure caption, accordingly.

By incorporating these intermediate dummy neuron populations, the overall delay in the network could be adjusted appropriately. Note that the rationale behind introducing dummy neurons into the network model was to shift the spike time of the presynaptic neuron by an amount determined mainly by the given axonal delay, while the weight of its (only) incoming synapse was of no importance, having no biological counterpart.

## Author Contributions

Conceptualization: A.V., A.K., A.A.; Methodology: A.G., H.R., A.V.; Investigation: A.G., H.R.; Supervision: A.V., A.K., A.A.; Writing – original draft: A.V.; Writing – review, and editing: A.K., A.A., A.V.

## References

[1] J. H. Kaas. Topographic maps are fundamental to sensory processing. Brain Research Bulletin 44.2 (1997), 107–112.

[2] A. Kohn, A. I. Jasper, J. D. Semedo, E. Gokcen, C. K. Machens, and B. M. Yu. Principles of corticocortical communication: proposed schemes and design considerations. Trends in Neurosciences 43.9 (2020), 725–737.

[3] P. Fries. A mechanism for cognitive dynamics: neuronal communication through neuronal coherence. Trends in Cognitive Sciences 9.10 (2005), 474–480.

[4] T. Akam and D. M. Kullmann. Oscillatory multiplexing of population codes for selective communication in the mammalian brain. Nature Reviews Neuroscience 15.2 (2014), 111–122.

[5] S. W. Wang and L. H. Tang. Emergence of collective oscillations in adaptive cells. Nature Communications 10.1 (2019), 5613.

[6] C. M. Gray, P. König, A. K. Engel, and W. Singer. Oscillatory responses in cat visual cortex exhibit inter-columnar synchronization which reflects global stimulus properties. Nature 338.6213 (1989), 334–337.

[7] A. Brovelli, M. Ding, A. Ledberg, Y. Chen, R. Nakamura, and S. L. Bressler. Beta oscillations in a large-scale sensorimotor cortical network: directional influences revealed by Granger causality. Proceedings of the National Academy of Sciences 101.26 (2004), 9849–9854.

[8] M. Bonnefond, S. Kastner, and O. Jensen. Communication between brain areas based on nested oscillations. eNeuro 4.2 (2017).

[9] A. M. Bastos, J. Vezoli, C. A. Bosman, J.-M. Schoffelen, R. Oostenveld, J. R. Dowdall, P. De Weerd, H. Kennedy, and P. Fries. Visual areas exert feedforward and feedback influences through distinct frequency channels. Neuron 85.2 (2015), 390–401.

[10] P. Fries. Neuronal gamma-band synchronization as a fundamental process in cortical computation. Annual Review of Neuroscience 32.1 (2009), 209–224.

[11] P. Fries. Rhythms for cognition: communication through coherence. Neuron 88.1 (2015), 220–235.

[12] T. Womelsdorf and P. Fries. The role of neuronal synchronization in selective attention. Current Opinion in Neurobiology 17.2 (2007), 154–160.

[13] F. Pouille and M. Scanziani. Enforcement of temporal fidelity in pyramidal cells by somatic feed-forward inhibition. Science 293.5532 (2001), 1159–1163.

[14] P. Fries, D. Nikolić, and W. Singer. The gamma cycle. Trends in Neurosciences 30.7 (2007), 309–316.

[15] A. Aertsen, M. Diesmann, and M.-O. Gewaltig. Propagation of synchronous spiking activity in feedforward neural networks. Journal of Physiology-Paris 90.3-4 (1996), 243–247.

[16] C. Haenschel, R. A. Bittner, J. Waltz, F. Haertling, M. Wibral, W. Singer, D. E. J. Linden, and E. Rodriguez. Cortical oscillatory activity is critical for working memory as revealed by deficits in early-onset schizophrenia. Journal of Neuroscience 29.30 (2009), 9481–9489.

[17] S. Neuenschwander and F. J. Varela. Visually triggered neuronal oscillations in the pigeon: an autocorrelation study of tectal activity. European Journal of Neuroscience 5.7 (1993), 870–881.

[18] G. J. Kress, M. J. Dowling, J. P. Meeks, and S. Mennerick. High threshold, proximal initiation, and slow conduction velocity of action potentials in dentate granule neuron mossy fibers. Journal of Neurophysiology 100.1 (2008), 281–291.

[19] K. Kusano. Electrical activity and structural correlates of giant nerve fibers in Kuruma shrimp (Penaeus japonicus). Journal of Cellular Physiology 68.3 (1966), 361–383.

[20] T. Pérez, G. C. Garcia, V. M. Eguíluz, R. Vicente, G. Pipa, and C. Mirasso. Effect of the topology and delayed interactions in neuronal networks synchronization. PLoS One 6.5 (2011), e19900.

[21] A. Pariz, I. Fischer, A. Valizadeh, and C. Mirasso. Transmission delays and frequency detuning can regulate information flow between brain regions. PLoS Computational Biology 17.4 (2021), e1008129.

[22] A. Ziaeemehr, M. Zarei, A. Valizadeh, and C. Mirasso. Frequency-dependent organization of the brain’s functional network through delayed-interactions. Neural Networks 132 (2020), 155–165.

[23] A. Knoblauch and G. Palm. Scene segmentation by spike synchronization in reciprocally connected visual areas. II. Global assemblies and synchronization on larger space and time scales. Biological Cybernetics 87.3 (2002), 168–184.

[24] I. Sugihara, E. J. Lang, and R. Llinás. Uniform olivocerebellar conduction time underlies Purkinje cell complex spike synchronicity in the rat cerebellum. The Journal of Physiology 470.1 (1993), 243–271.

[25] O. C. Time. Role of myelination in the development of a uniform olivocerebellar conduction. Journal of Neurophysiology 89 (2003), 2259–2270.

[26] M. Salami, C. Itami, T. Tsumoto, and F. Kimura. Change of conduction velocity by regional myelination yields constant latency irrespective of distance between thalamus and cortex. Proceedings of the National Academy of Sciences 100.10 (2003), 6174–6179.

[27] J. G. Pelletier and D. Paré. Uniform range of conduction times from the lateral amygdala to distributed perirhinal sites. Journal of Neurophysiology 87.3 (2002), 1213–1221.

[28] T. Chomiak, S. Peters, and B. Hu. Functional architecture and spike timing properties of corticofugal projections from rat ventral temporal cortex. Journal of Neurophysiology 100.1 (2008), 327–335.

[29] A. H. Seidl. Regulation of conduction time along axons. Neuroscience 276 (2014), 126–134.

[30] S. Pajevic, D. Plenz, P. J. Basser, and R. D. Fields. Oligodendrocyte-mediated myelin plasticity and its role in neural synchronization. eLife 12 (2023), e81982.

[31] R. D. Fields. White matter in learning, cognition and psychiatric disorders. Trends in Neurosciences 31.7 (2008), 361–370.

[32] J. M. Williamson and D. A. Lyons. Myelin dynamics throughout life: an ever-changing landscape? Frontiers in Cellular Neuroscience 12 (2018), 424.

[33] W. Gerstner, R. Kempter, J. L. Van Hemmen, and H. Wagner. A neuronal learning rule for sub-millisecond temporal coding. Nature 383.6595 (1996), 76–78.

[34] R. R. Kerr, A. N. Burkitt, D. A. Thomas, M. Gilson, and D. B. Grayden. Delay selection by spike-timing-dependent plasticity in recurrent networks of spiking neurons receiving oscillatory inputs. PLoS Computational Biology 9.2 (2013), e1002897.

[35] G. Hahn, A. F. Bujan, Y. Frégnac, A. Aertsen, and A. Kumar. Communication through resonance in spiking neuronal networks. PLoS Computational Biology 10.8 (2014), e1003811.

[36] H. Rezaei, A. Aertsen, A. Kumar, and A. Valizadeh. Facilitating the propagation of spiking activity in feedforward networks by including feedback. PLoS Computational Biology 16.8 (2020), e1008033.

[37] B. Katz and R. Miledi. The measurement of synaptic delay, and the time course of acetylcholine release at the neuromuscular junction. Proceedings of the Royal Society of London. Series B. Biological Sciences 161.985 (1965), 483–495.

[38] L. Stanford. Conduction velocity variations minimize conduction time differences among retinal ganglion cell axons. Science 238.4825 (1987), 358–360.

[39] S. Boudkkazi and D. Debanne. Enhanced release probability without changes in synaptic delay during analogue–digital facilitation. Cells 13.7 (2024), 573.

[40] S. Boudkkazi, E. Carlier, N. Ankri, O. Caillard, P. Giraud, L. Fronzaroli-Molinieres, and D. Debanne. Release-dependent variations in synaptic latency: a putative code for short- and long-term synaptic dynamics. Neuron 56.6 (2007), 1048–1060.

[41] S. Boudkkazi, L. Fronzaroli-Molinieres, and D. Debanne. Presynaptic action potential waveform determines cortical synaptic latency. The Journal of Physiology 589.5 (2011), 1117–1131.

[42] D. J. Bakkum, Z. C. Chao, and S. M. Potter. Long-term activity-dependent plasticity of action potential propagation delay and amplitude in cortical networks. PLOS One 3.5 (2008), e2088.

[43] Q. Yu, J. Gao, J. Wei, J. Li, K. C. Tan, and T. Huang. Improving multispike learning with plastic synaptic delays. IEEE Transactions on Neural Networks and Learning Systems 34.12 (2022), 10254–10265.

[44] R. M. Wang, T. J. Hamilton, J. C. Tapson, and A. van Schaik. A neuromorphic implementation of multiple spike-timing synaptic plasticity rules for large-scale neural networks. Frontiers in Neuroscience 9 (2015), 180.

[45] E. M. Izhikevich. Polychronization: computation with spikes. Neural Computation 18.2 (2006), 245–282.

[46] L. C. Katz and C. J. Shatz. Synaptic activity and the construction of cortical circuits. Science 274.5290 (1996), 1133–1138.

[47] S. LeVay and M. P. Stryker. “The development of ocular dominance columns in the cat”. Soc. Neurosci. Symp. Vol. 4. 1979, 83–98.

[48] R. P. Rao and D. H. Ballard. Predictive coding in the visual cortex: a functional interpretation of some extra-classical receptive-field effects. Nature Neuroscience 2.1 (1999), 79–87.

[49] D. H. Hubel, T. N. Wiesel, et al. Receptive fields of single neurones in the cat’s striate cortex. J Physiol 148.3 (1959), 574–591.

[50] T. K. Hensch. Critical period plasticity in local cortical circuits. Nature Reviews Neuroscience 6.11 (2005), 877–888.

[51] P. J. Uhlhaas and W. Singer. Abnormal neural oscillations and synchrony in schizophrenia. Nature Reviews Neuroscience 11.2 (2010), 100–113.

[52] A. Kumar and M. R. Mehta. Frequency-dependent changes in NMDAR-dependent synaptic plasticity. Frontiers in Computational Neuroscience 5 (2011), 38.

[53] W. Gerstner. Time structure of the activity in neural network models. Physical Review E 51.1 (1995), 738.

[54] A. Sahasranamam, I. Vlachos, A. Aertsen, and A. Kumar. Dynamical state of the network determines the efficacy of single neuron properties in shaping the network activity. Scientific Reports 6.1 (2016), 26029.

[55] F. Devalle and A. Roxin. How plasticity shapes the formation of neuronal assemblies driven by oscillatory and stochastic inputs. Journal of Computational Neuroscience 53.1 (2025), 9–23.

[56] G. Q. Bi and M. M. Poo. Synaptic modifications in cultured hippocampal neurons: dependence on spike timing, synaptic strength, and postsynaptic cell type. Journal of Neuroscience 18.24 (1998), 10464–10472.

[57] H. X. Wang, R. C. Gerkin, D. W. Nauen, and G. Q. Bi. Coactivation and timingdependent integration of synaptic potentiation and depression. Nature Neuroscience 8.2 (2005), 187–193.

[58] W. M. Kistler and J. L. van Hemmen. Modeling synaptic plasticity in conjunction with the timing of pre-and postsynaptic action potentials. Neural Computation 12.2 (2000), 385–405.

[59] R. C. Froemke and Y. Dan. Spike-timing-dependent synaptic modification induced by natural spike trains. Nature 416.6879 (2002), 433–438.

[60] E. L. Bienenstock, L. N. Cooper, and P. W. Munro. Theory for the development of neuron selectivity: orientation specificity and binocular interaction in visual cortex. Journal of Neuroscience 2.1 (1982), 32–48.

[61] S. Song, K. D. Miller, and L. F. Abbott. Competitive Hebbian learning through spiketiming-dependent synaptic plasticity. Nature Neuroscience 3.9 (2000), 919–926.

[62] C. Clopath, L. Büsing, E. Vasilaki, and W. Gerstner. Connectivity reflects coding: a model of voltage-based STDP with homeostasis. Nature Neuroscience 13.3 (2010), 344–352.

[63] J. Rubin, D. D. Lee, and H. Sompolinsky. Equilibrium properties of temporally asymmetric Hebbian plasticity. Physical Review Letters 86.2 (2001), 364.

[64] M. C. Van Rossum, G. Q. Bi, and G. G. Turrigiano. Stable Hebbian learning from spike timing-dependent plasticity. Journal of Neuroscience 20.23 (2000), 8812–8821.

[65] E. J. Agnes and T. P. Vogels. Co-dependent excitatory and inhibitory plasticity accounts for quick, stable and long-lasting memories in biological networks. Nature Neuroscience 27.5 (2024), 964–974.

[66] D. E. Feldman. The spike-timing dependence of plasticity. Neuron 75.4 (2012), 556–571.

[67] X. J. Wang. Neurophysiological and computational principles of cortical rhythms in cognition. Physiological Reviews 90.3 (2010), 1195–1268.

[68] N. Brunel. Dynamics of sparsely connected networks of excitatory and inhibitory spiking neurons. Journal of Computational Neuroscience 8.3 (2000), 183–208.

[69] S. Toth, A. Solyom, J. Vajda, and Z. Toth. The rhythmic properties of the motor system. Stereotactic and Functional Neurosurgery 53.2 (1989), 95–104.

[70] A. Kumar, S. Rotter, and A. Aertsen. Spiking activity propagation in neuronal networks: reconciling different perspectives on neural coding. Nature Reviews Neuroscience 11.9 (2010), 615–627.

[71] M.-O. Gewaltig and M. Diesmann. Nest (neural simulation tool). Scholarpedia 2.4 (2007), 1430.

[72] C. Linssen and et al. NEST 2.16.0. 2018. doi: 10.5281/zenodo.1400175. URL: https://doi.org/10.5281/zenodo.1400175.

[73] A. Morrison, S. Straube, H. E. Plesser, and M. Diesmann. Exact subthreshold integration with continuous spike times in discrete-time neural network simulations. Neural Computation 19.1 (2007), 47–79.

